# Spatial dissection of a megabase-scale gene in *Drosophila* spermatocyte nuclei

**DOI:** 10.64898/2026.11.22.768416

**Authors:** McKenna Elise Rook, E.K. Beard, Fumihiro Watanabe, Mayu Inaba

## Abstract

Understanding how nascent RNA is structurally organized during transcription remains challenging for most genes because their transcription units are too small to resolve the spatial organization of nascent transcripts. Exceptionally long genes provide a unique opportunity to address how RNA is organized during transcription. Here, we investigate the transcriptional organization of *kl-3*, an exceptionally long Y-linked fertility gene expressed in *Drosophila* spermatocytes. Using RNA FISH against AATAC satellite sequence embedded in a large intron (intron5), we find that nascent exon-containing transcripts remain tightly associated with the Y-loop DNA strand, whereas intronic transcripts form a distinct outer layer surrounding the exon-associated signal. We further demonstrate that Smooth protein associates specifically with the AATAC positive loop, and is required for proper transcript organization, likely by creating phase-separated compartments. Finally, we identify distinct subnuclear localization of transcription-, elongation-, and splicing-machineries. H3K4me3 is enriched near the nucleolus-like body (NLB), consistent with transcriptional initiation in this region. Unexpectedly, we found the splicing-associated domain, H3K36me3, and splicing factors including U2af38 and Cdc5, are also enriched near the NLB, suggesting that splicing may be spatially undertaken within this compartment. Together, our findings reveal a spatial organization of nascent RNA in which DNA-associated exon sequences and compacted intronic transcripts are coordinated with distinct transcriptional and RNA-processing subnuclear domains. We propose that such spatial segregation may be commonly utilized in broad cellular genes.

## Introduction

Eukaryotic pre-mRNA splicing often occurs co-transcriptionally, requiring precise coordination between transcription elongation, RNA folding, chromatin organization, and spliceosome assembly. Spliceosome assembly is initiated by the recruitment of U1 snRNP to the 5′ splice site of nascent pre-mRNA, followed by the recruitment of U2 snRNP to the branch-point sequence, forming the prespliceosome. Then the U4/U6.U5 tri-snRNP is recruited and catalytic remodeling occurs to excise the intron via two sequential transesterification reactions (reviewed in ^1^). Pre-mRNA splicing has traditionally been understood through structural studies in which spliceosomal components recognize and assemble on specific sequences within pre-mRNA. However, because a substantial fraction of pre-mRNA splicing occurs co-transcriptionally, a fundamental question remains: how is spliceosome assembly coordinated with the transcriptional machinery on nascent RNA, and how are these functional components organized on the template DNA strand in vivo? Multiple studies have determined molecular interaction between splicing and transcriptional components (reviewed in ^2^). Moreover, recent studies using cryo-electron microscopy has provided the structure of a transcribing RNA polymerase II (Pol II)–U1 snRNP complex which revealed direct interactions between Pol II and U1 snRNP, positioning the nascent 5′ splice site near the RNA exit channel of Pol II ^3^. As transcription proceeds, the retained 5′ splice site forms a growing intron loop, which facilitates scanning of nascent pre-mRNA for the downstream 3′ splice site and promotes efficient spliceosome assembly ^3,4^. These findings support a model in which spliceosome assembly is physically coordinated with transcription elongation through interactions involving the Pol II C-terminal domain and splicing factors, including U2AF and U2 snRNP ^5^.

Despite these advances, examining how these molecular interactions are organized and coordinated in vivo remains challenging. The relatively small dimensions of transcription and splicing complexes, together with their dynamic and transient interactions within the nucleus, make it difficult to resolve their spatial organization and capture their functional relationships in intact cells using conventional imaging approaches. Recent advances in single-molecule RNA imaging have begun to provide insights into dynamic processes underlying co-transcriptional splicing ^6–9^. How these molecular interactions and regulatory events are spatially organized within the nucleus in vivo, however, remains poorly understood.

The Y chromosome fertility genes of *Drosophila melanogaster* provide a unique opportunity to overcome this limitation. These genes contain giant megabase-scale introns and are transcribed during spermatocyte development to produce characteristic lampbrush chromosome loops, or Y-loops. Each loop (Y-loop A, B, or C) contains a single large fertility gene (*kl-5*, *kl-3* and *ORY/ks-1*, respectively) on which nascent transcripts remain associated with their sites of transcription while undergoing RNA processing ^10–14^. Because these loops represent some of the largest naturally occurring transcription units, they have served as classical models for studying transcriptional organization for decades ^15^. Structural studies of the Y-loops have revealed a multilayered architecture comprising distinct RNA- and protein-rich domains ^16,17^.

Recent work by Fingerhut and colleagues demonstrated that transcription proceeds in a highly coordinated spatiotemporal manner throughout spermatocyte differentiation, which spans approximately 90 hours ^18^. Transcription initiates at the 5′ regions in early spermatocytes, followed by transcription of intronic satellite DNA sequences, whereas distal exons are transcribed only during the later stages of spermatocyte development. Mature transcripts subsequently accumulate in the cytoplasm as distinct RNA granules ^18^. More recently, Fingerhut et al. demonstrated that co-transcriptional splicing is required for efficient transcription of these gigantic genes ^19^. Depletion of splicing factors resulted in transcriptional attenuation within giant introns, accompanied by disorganization and intermingling of nascent RNA domains ^19^. These findings suggest the importance of the structural regulation of this process.

Despite these advances in understanding the temporal progression of transcription and the functional requirement for co-transcriptional splicing, the spatial organization of nascent transcripts remains poorly understood. In particular, the repetitive nature of giant introns has prevented visualization of individual intronic regions, limiting mechanistic investigation of how intronic sequences are organized.

Our previous study has overcome this long-standing barrier by identifying transposable element (TE) insertions embedded within the giant introns of multiple Y-linked fertility genes ^20^. These intronic TEs serve as endogenous molecular landmarks that enable visualization of discrete intronic regions during transcription with unprecedented detail ^20^. In this study, we identified two structural principles of *kl-3* gene regulation. First, we reveal an unexpected higher-order organization of nascent RNA in which exon sequences remain tightly associated with the DNA template strand, while intronic RNA is compactly organized as an outer layer surrounding the exon-associated RNA. We demonstrate that this intronic RNA compaction is regulated by the RNA-binding protein Smooth, and that disruption of this organization impairs elongation of the gene towards later exons. Second, we show that key splicing components involved in 3′ splice-site recognition, localizes near the nucleolus-like body (NLB), a region recently implicated in Pol II activation and transcriptional initiation by Fingerhut et al. (bioRxiv, 2026)^21^. This localization suggests that transcriptional initiation and spliceosome recruitment may be spatially coordinated near the NLB, suggesting a spatially compartmentalized chromatin organization, potentially reflecting the positioning of exon–intron boundaries near the NLB and intronic regions along the loops.

Together, our findings establish a previously unrecognized intron-loop architecture that spatially organizes nascent RNA and facilitate transcription. Moreover, our data suggest the special segregation of distinct processes of gene expression within a same gene locus.

## Results

### Nascent kl-3 transcripts associate tightly with the DNA template strand

To investigate the spatial organization of nascent transcripts on a Y chromosome fertility gene, *kl-3*, we performed RNA fluorescence in situ hybridization (FISH) to visualize nascent transcripts throughout the stage progression of spermatocytes (SCs) in the *Drosophila* testes (Figure 1A). In SCs, Y-loops are known to spread out in the nucleoplasm surrounded by chromosome territories associated with nuclear periphery (Figure 1B). Using the probe targeting *kl-3* exon 1, we observed a characteristic loop pattern along the DAPI positive Y-loops (Figure 1C).

**Figure 1.**
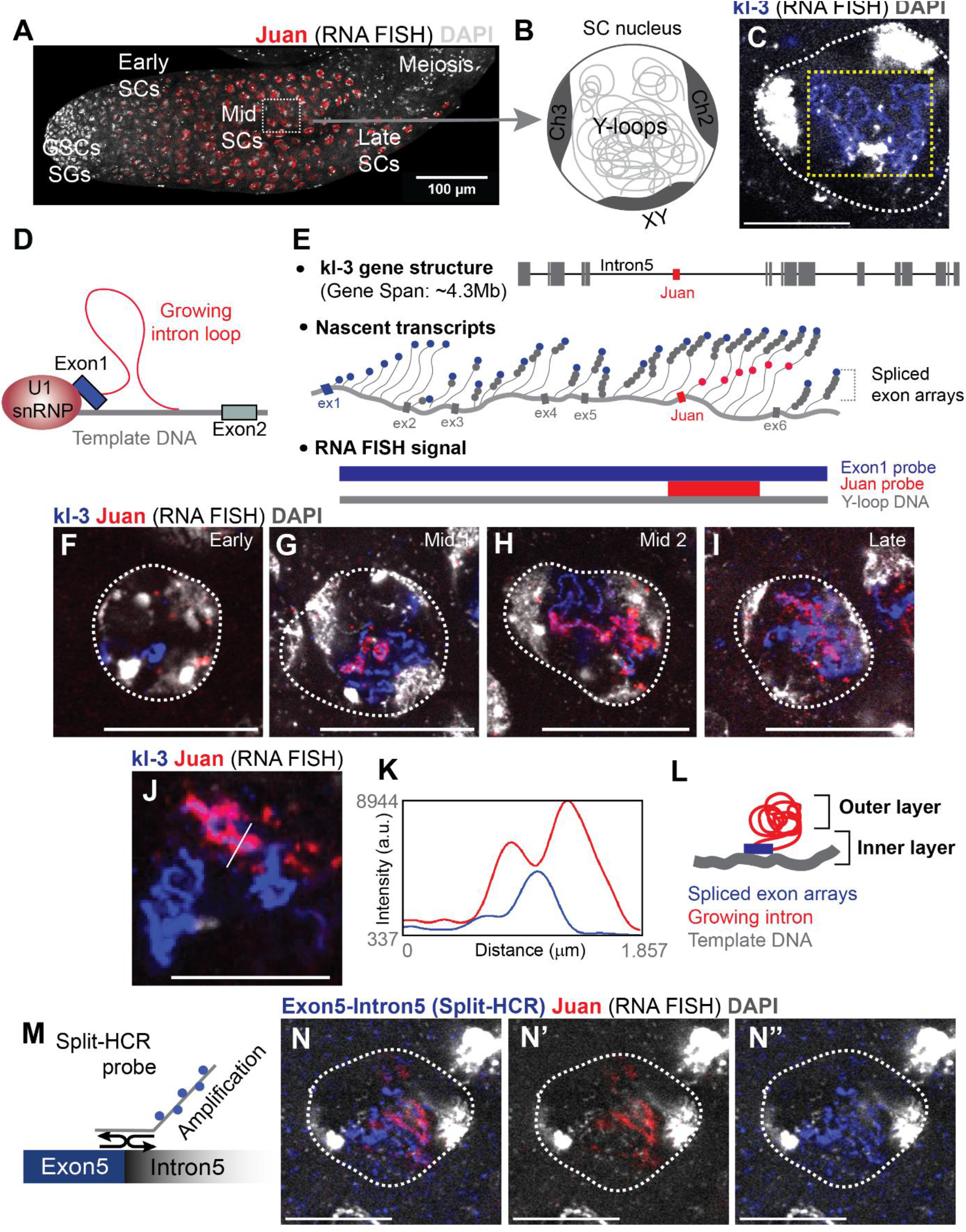
**(A)** Representative RNA FISH image of a testis containing developing germ cells, including germline stem cells (GSCs), spermatogonia (SGs), and early-, mid-, and late-stage spermatocytes (SCs). RNA FISH targeting the Juan transposable element reveals strong signals in the nucleoplasm of spermatocytes (red). DAPI (gray). **(B)** Schematic representation of a *Drosophila* SC nucleus. Chromosomes occupy distinct territories, with specific chromatin positioning and associations with the nuclear envelope. In the nuclear interior, Y-linked fertility genes form giant, individually resolvable chromatin loops (e.g., Y-loops A, B, and C). **(C)** Representative RNA FISH image of a mid-stage SC using a probe against the *kl-3* exon 1 sequence (blue). DAPI (gray) highlights the Y-loop chromatin fiber underlying the *kl-3* signal. A white dotted line outlines the SC nucleus. A yellow dotted rectangle outlines the *kl-3*-positive Y-loop. **(D)** Schematic representation of the growing intron loop model. **(E)** Schematic representations of organization of the *kl-3* gene products, the composition of the nascent transcript, and the predicted RNA FISH signal distribution along the loop. **(F–I)** Representative RNA FISH images of early- (F), mid- (G and H), and late-stage (I) SCs using a probe against the *kl-3* exon 1 sequence (blue) and a Juan probe (red). DAPI (gray). White dotted lines outline SC nuclei. **(J–K)** Representative RNA FISH image of a mid-stage SC using a probe against the *kl-3* exon 1 sequence (blue) and a Juan probe (red). The white line indicates the region used for intensity measurements, as shown in the line plot in (K). DAPI (gray). **(L)** Schematic interpretation of the observed two-layer structure. **(M)** Schematic representation of the Split-HCR probe designed to detect the exon5-intron5 junction. **(N–N″)** Representative RNA FISH images of a mid-stage SC using probes against Juan (red) and the exon5-intron5 junction (Split-HCR, blue). White dotted lines outline SC nuclei. All scale bars are 10 µm unless otherwise indicated in the images.

The close association between the exon signal and the DNA strand well reflects a previously proposed model, the "growing intron loop” model, in which spliced 5’ exons and 5’ splicing factors are located close to the newly synthesized nascent pre-mRNA to scan for the 3’ splice site (Figure 1D) ^3,4^. To further confirm this model, we attempted to resolve the relative localization of the exon and intron sequences within nascent transcripts. However, visualizing individual introns within these giant Y-loop genes has been challenging, mainly because of their highly repetitive sequences, which remain largely unresolved at the genomic level ^22^. Our previous study has suggested that several transposable elements (TEs) are embedded within giant introns of Y-loop genes, and the FISH probes against these TEs enable targeting of specific giant intron ^20^. Therefore, we used the probes against two TEs, *Juan* and *HMS-Beagle*, located within introns of the *kl-3* and *kl-5* genes, respectively to overcome the issue ^20^. The *Juan* insertion is located within intron5, an extremely long intron in the *kl-3* gene (Figure 1E) ^20^.

The size of *kl-3* positive area gradually increases as SC’s stage progresses, consistent with a previous report ^18^ (Figure 1F-I). As we reported previously ^20^, *Juan* signals emerge closely associating with a part of the *kl-3* positive loop, starting from mid-stage SCs and extending toward late-stage SCs (Figure 1G-I). In late-stage SCs, we also observed dotty *Juan* signals surrounding the *kl-3* positive loop, likely reflecting the spliced fraction of the intron5 (Figure 1I).

Next, we performed closer observation of the *kl-3* positive loop to assess the spatial relationship between the *Juan* positive intronic signal and the *kl-3* exon1 signal. In particular, *Juan* signals appeared to extend outside the tightly DNA-associated exon1 signal (Figure 1J, K), suggesting that intronic RNA sequences occupy a spatially distinct layer surrounding the exon-associated RNA. This organization is consistent with a “growing intron loop”-model, and this further suggests that intronic sequences, which likely grow beyond megabase size, are packed into a relatively compact size, to form a tightly aligned outer layer to the exon-containing transcripts (Figure 1L).

We also examined *HMS-Beagle*, another transposable element localized in a kl-5 intron, another Y-loop forming gene. We observed a similar two-layer pattern, such that the *kl-5* exon probe signal is closely associated with the template DNA, while intron RNA visualized by the *HMS-Beagle* probe illuminates an outer layer (Figure S1A), suggesting the possible commonality of the spatial organization of nascent transcripts.

To confirm that the intronic signal throughout the Y-loop is truly unspliced nascent transcripts, we performed FISH that detects only the *kl-3* exon-intron (exon5-intron5) boundary by using two short split HCR (Hybridization chain reaction) probes, one which binds to the end of exon5 and the other which binds to the start of intron5, and both probes are necessary to be adjacent to start signal amplification (Figure 1M). As we expected, we observed HCR FISH signal overlaps with the *Juan* positive loop, closely associated with the DNA strand (Figure 1N), confirming that the detected *Juan* is part of unspliced nascent transcript (Figure 1E).

Together, these observations reveal a distinct spatial organization of nascent transcripts within the Y-loop, in which exon-associated sequences closely follow the DNA template strand, while intronic sequences form a separate, compacted outer layer (Figure 1L). This organization is consistent with the previously proposed “growing intron loop” model ^3^, suggesting a possible relevance for functional coupling of transcription and splicing ^3^.

### Intronic transcripts are folded as they elongate on distal Y-loops

During transcription, nascent RNA is folded into secondary or higher structures as it emerges from RNA polymerase^23,24^. Nascent RNA folding can influence co-transcriptional RNA processing, including splicing efficiency and splice-site selection, suggesting that RNA structure is closely linked to the regulation of gene expression ^25–28^. To further investigate how *kl-3* intronic transcripts are organized along the extended Y-loop structure, we sought to identify satellite DNA sequences that could cover a larger length of intron5. The AATAT satellite sequence is known to be embedded within multiple *kl-3* introns as well as other genomic regions ^29^. Therefore, AATAT cannot serve as a specific marker for an individual *kl-3* intron. We found that the AATAC satellite repeat blocks, which was originally mapped around the *kl-3* locus, is located closely to one of the AATAT repeats in Y chromosomes (Figure 2A). This localization is consistent with a previous study mapping satellite repeats across *Drosophila* strains, which identified satellite arrays exceeding 100 Kb, suggesting that the repeat region spans more than 100 Kb ^30^. DNA FISH using an AATAC probe illuminates a part of Y-loop structure in SCs, indicating that AATAC repeats are likely located within the *kl-3* Y-loop (Figure 2B, C). Indeed, RNA-FISH against AATAC shows perfect colocalization with *Juan* on the intronic loop region (Figure 2D). In combination with cytological mapping of AATAC repeat localization around *kl-3*, these observations suggest that AATAC sequences are embedded within intron 5 of *kl-3* gene, likely close to- or after- the *Juan* insertion site. Thus, AATAC repeats provide an additional marker for examining the spatial distribution of intronic sequences along the *kl-3* loop.

**Figure 2.**
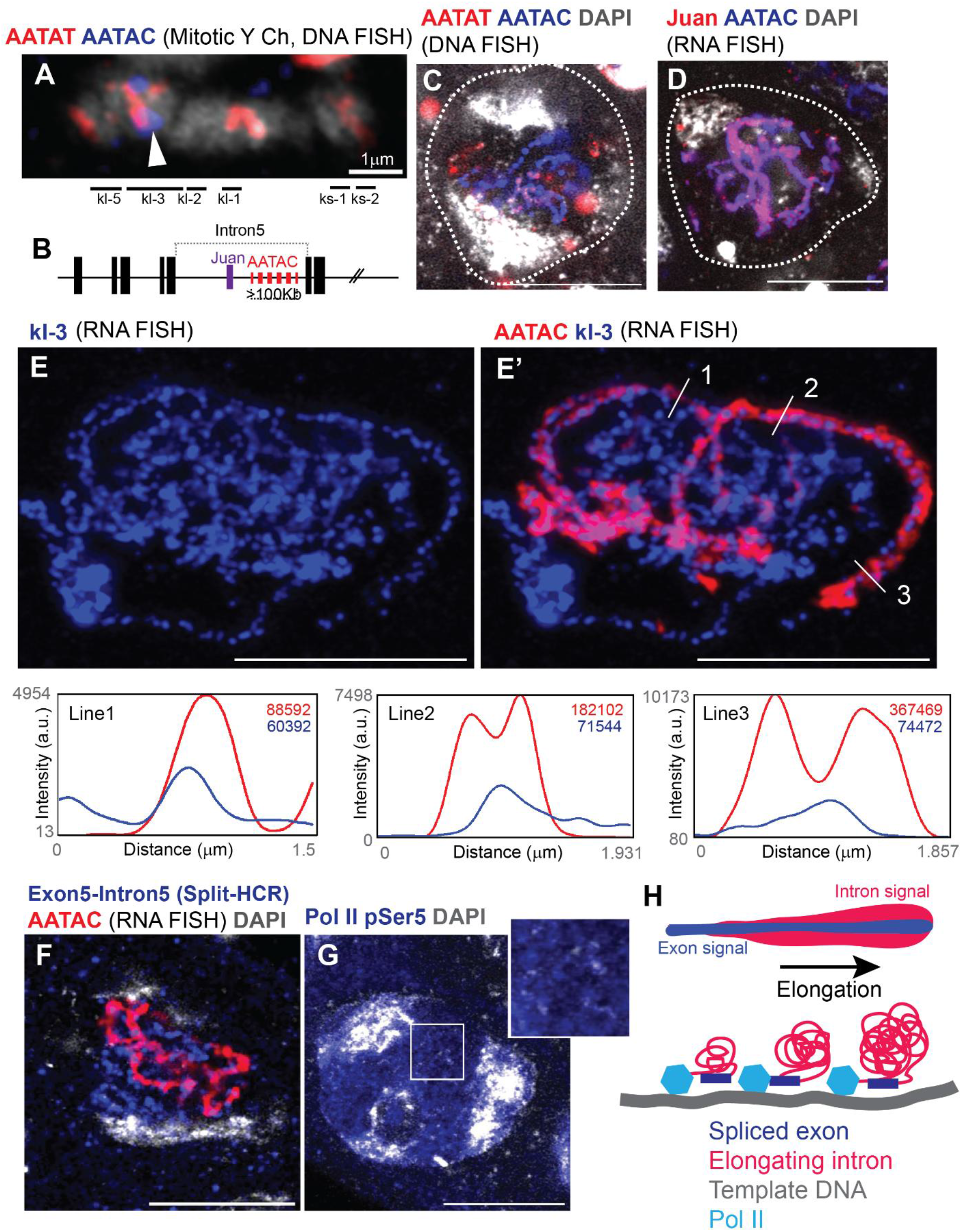
**(A)** DNA FISH analysis of the Y chromosome in neuroblast chromosome spreads from *w¹¹¹⁸* flies. The approximate locations of major Y-linked fertility genes are indicated based on Bonaccorsi et al.^10^ The probes used are shown in the corresponding colors. The arrowhead indicates an AATAC signal within the *kl-3* locus. **(B)** Predicted size and localization of AATAC satellite blocks within *kl-3* intron5. **(C)** Representative DNA FISH image of SC nuclei from *w¹¹¹⁸* flies. The probes used are shown in the corresponding colors. White dotted lines outline SC nuclei. **(D)** Representative RNA FISH image of SC nuclei from *w¹¹¹⁸* flies. The probes used are shown in the corresponding colors. White dotted lines outline SC nuclei. **(E–E′)** Representative RNA FISH images of SC nuclei from *w¹¹¹⁸* flies. The probes used are shown in the corresponding colors. White lines indicate the regions used for the corresponding line scans shown in the lower panels. The line scans show fluorescence intensity along the indicated distances. Colored values indicate the integrated fluorescence intensity for each channel across the scanned region. **(F)** Representative RNA FISH image of an SC nucleus from *w¹¹¹⁸* flies. The probes used are shown in the corresponding colors. White dotted lines outline the SC nucleus. **(G)** Representative immunofluorescence image of an SC nucleus from *w¹¹¹⁸* flies stained with an antibody against Pol II pSer5. The boxed region is magnified in the upper-right panel. **(H)** Schematic interpretation of the elongating *kl-3* loop, illustrating the specific organization of spliced exon arrays and growing introns. DAPI (gray). Scale bars are 10 µm unless otherwise indicated in the images.

RNA FISH using probes against the AATAC sequence revealed intronic signals extending along the template Y-loop, with the signal progressively increasing in thickness along its length (Figure 2E). This gradual increase in AATAC signal intensity suggests that intron5 contains a long AATAC array or multiple repeat blocks (Figure 2B). The progressive expansion of the intronic signal along the Y-loop is also consistent with ongoing intron elongation accompanied by folding of the nascent transcript. Notably, the intronic signal extended to approximately 1 μm in thickness (Figure 2E, line scans), suggesting extensive compaction and folding of the nascent intron are occurring during elongation. Split-HCR to detect the exon5-intron5 boundary revealed that AATAC is indeed located in intron5 (Figure 2F). Consistent with possible active elongation is occurring throughout the loops, we detected phosphorylated RNA polymerase II at Ser5 (Pol II-pSer5) spread throughout the entire nucleoplasm, providing evidence that transcriptionally engaged polymerase is present beyond the proximal region (Figure 2G).

Together, the detection of intronic RNA throughout the Y-loop, combined with the presence of phosphorylated RNA polymerase II, supports a model in which transcription extends throughout the distal regions of the Y-loop. These findings further suggest that intronic transcripts are tightly packed and remain associated with the transcriptional structure over an extended distance (Figure 2H).

### Disruption of intronic transcript compaction impairs progression of *kl-3* transcription

A specialized organization of intron and exon regions within nascent transcripts led us to investigate whether the observed organization contributes to proper gene expression. Therefore, we sought to interfere with the organization of intronic sequences to see the effect on *kl-3* expression.

It has been known that Y-loops are associated with specific RNA binding proteins ^18,31–33^. Blanks has been shown to be concentrated around the *kl-3* Y-loop (Figure 3A) ^18^. By careful observation of Blanks-GFP pattern, we noticed that Blanks does not overlap with the AATAC-positive region but instead localizes to the outer boundary of the loop, surrounding the entire AATAC-positive domain (Figure 3A). Our further investigation of candidate RNA-binding proteins previously implicated in Y-loop localization^31^ revealed that Smooth (Sm) is enriched in close proximity to the AATAC-positive region of the *kl-3* Y-loop (Figure 3B). Sm-GFP is completely overlapped with the layered organization of nascent transcripts (Figure 3C, D).

**Figure 3.**
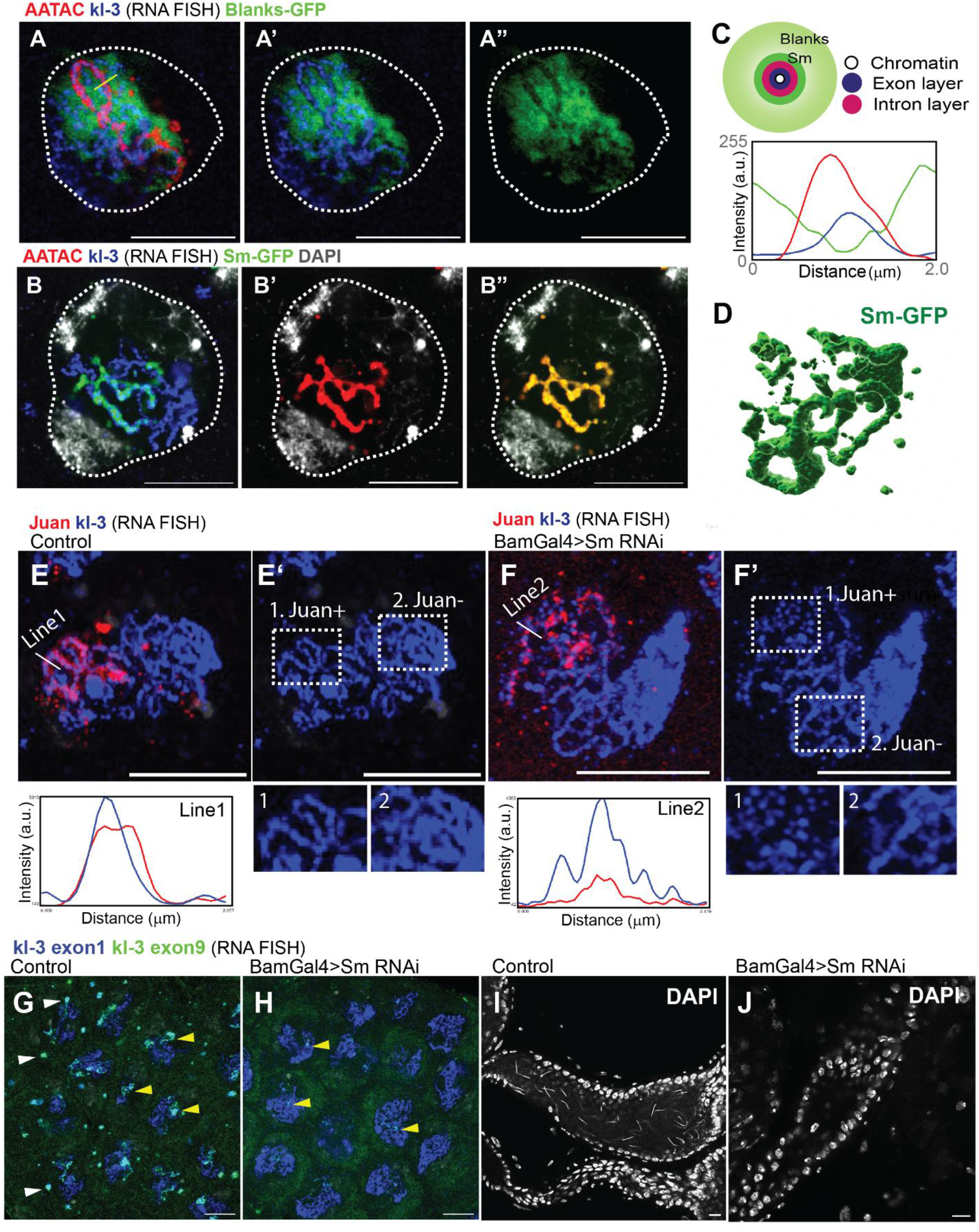
**(A, B)** Representative RNA FISH images of SC nuclei from Blanks-GFP (A) or Sm-GFP (B) flies. The probes used are shown in the corresponding colors. White dotted lines outline SC nuclei. The yellow line in (A) indicates the region used for the line scan shown in (C). **(C)** Interpretation of Blanks and Sm localization relative to elongating *kl-3* loops. The line scan corresponds to the yellow line indicated in (A). **(D)** Representative 3D rendering of Sm-GFP signals within SC nuclei. Images were analyzed using Imaris software. **(E, F)** Representative RNA FISH images of mid-stage SCs hybridized with the indicated probes in testes from control (*bamGal4*) flies (E) or *bamGal4*-driven *Sm RNAi* flies (F). The probes used are shown in the corresponding colors. White lines indicate the regions used for the corresponding line scans shown in the lower panels. The line scans show fluorescence intensity along the indicated distances. In (E′) and (F′), white dotted squares mark *Juan*-positive and *Juan*-negative *kl-3* loops, respectively, which are shown in the lower panels. In (F′), the *Juan*-positive loop, corresponding to a later stage of intron5 transcription, shows a disrupted layered structure. **(G, H)** Representative RNA FISH images of mid-stage SCs hybridized with the indicated probes in testes from control (*bamGal4*) flies (G) or *bamGal4*-driven *Sm RNAi* flies (H). White arrowheads in (G) indicate *kl-3* exon 9 FISH signals in cytoplasmic *kl* bodies, which are absent in *Sm RNAi* testes (H). Yellow arrowheads indicate exon 9-positive regions along the loops, corresponding to nascent transcripts associated with the 3′ side of the *kl-3* loop downstream of exon 9. **(I, J)** Representative DAPI-stained images of seminal vesicles (SVs) from flies of the indicated genotypes. Scale bars are 10 µm.

Striking co-localization of Sm-GFP with the AATAC containing intron prompted us to investigate whether Sm contributes to the formation and maintenance of a compartment along the *kl-3* Y-loop. We first examined whether the loss of either protein affected the organization of the layered organization of nascent transcripts. Indeed, Sm RNAi testes showed disruption of the two-layered organization of exon and intron, indicating that Sm is required to maintain their organization (Figure3E, F). The altered intronic RNA distribution was accompanied by a reduction or loss of the mature *kl-3* cytoplasmic RNA body, a distinct cytoplasmic structure enriched in mature *kl-3* transcripts (Figure3G, H). Moreover, consistent with the essential role of *kl-3* gene product encoding axonemal outer arm dynein heavy chain in male fertility, loss of mature *kl-3* transcripts resulted in lack of mature sperm in seminal vesicles (Figure 3I, J).

Together, these results indicate that Sm is required for the proper spatial organization of intronic transcripts and that this organization is associated with successful *kl-3* transcription and maturation. These findings suggest that layered organization of the exon array and intronic transcript is not simply a consequence of transcription but may actively contribute to the production of mature *kl-3* mRNA.

### Sm-GFP is fully mobile within compartments

We next wondered how Sm contributes to the formation and maintenance of a compartment along the *kl-3* Y-loop. Specifically, we wondered if Sm protein binds to nascent transcript to modulate the structure, or if the protein rather contributes to creating an environment via formation of a phase-separated compartment. To distinguish these possibilities, we performed fluorescence recovery after photobleaching (FRAP) to examine the molecular dynamics of Sm-GFP. Following photobleaching of GFP fluorescence in a defined region of the loop-associated compartment, we monitored fluorescence recovery over time. Interestingly, we observed rapid recovery of the photobleached region through the redistribution of Sm-GFP molecules from unbleached regions of the loop-associated compartment. This rapid recovery indicates that Sm-GFP molecules are highly mobile within the compartment and are not stably bound to pre-existing nascent intronic transcripts (Figure 4A-C, Movie S1).

**Figure 4.**
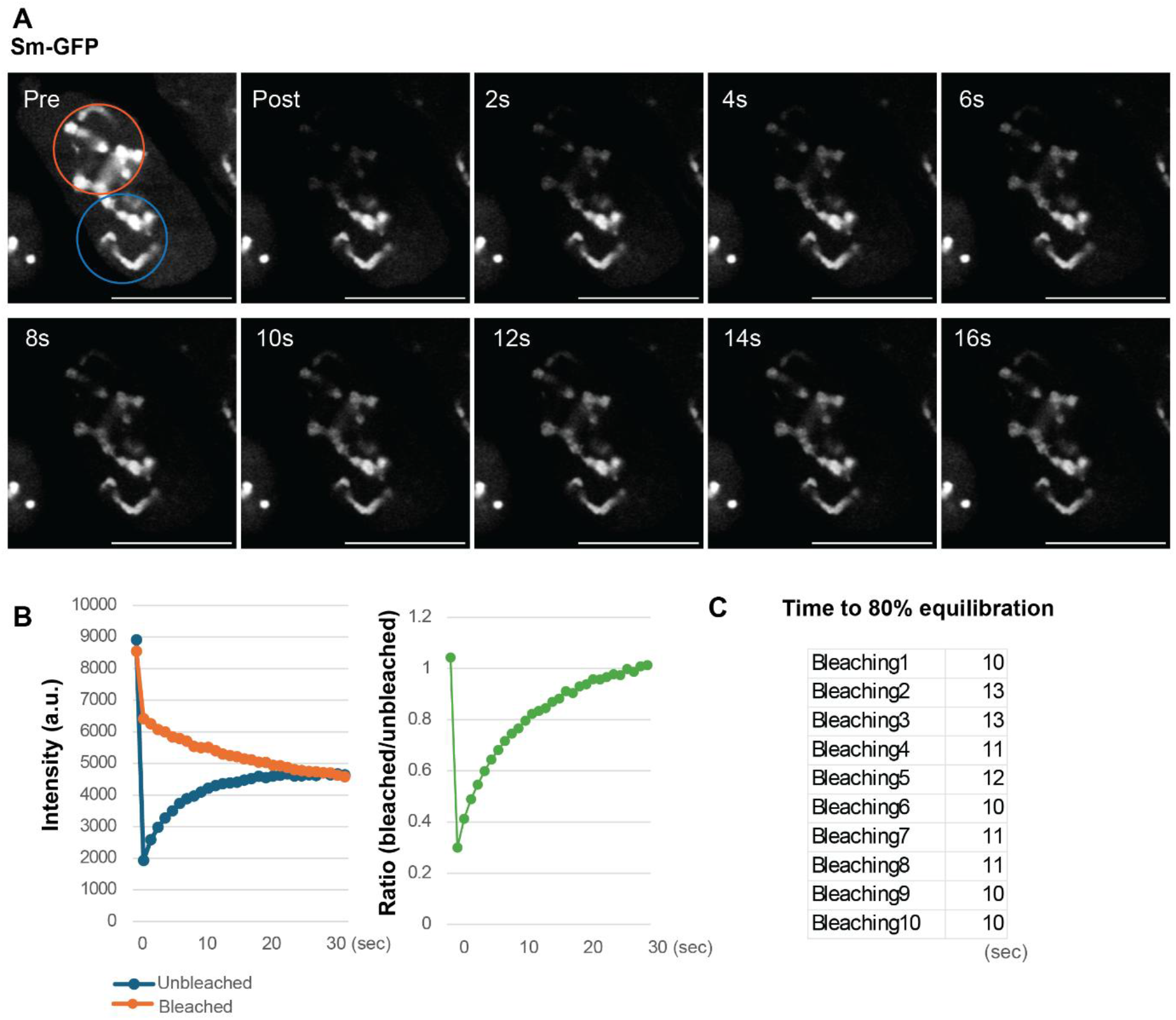
**(A)** Representative images from a Sm-GFP photobleaching experiment in a mid-stage SC expressing Sm-GFP. Images taken before photobleaching (pre-bleaching) and at the indicated time points (seconds) after photobleaching are shown. The orange line indicates the photobleached region (half of the nucleoplasm). The blue circle indicates the region used for intensity measurements. Sm-GFP fluorescence intensity rapidly equilibrates between the bleached and unbleached regions. **(B)** Measurement of Sm-GFP fluorescence intensities after photobleaching in the cell shown in (A). The left graph shows the measured fluorescence intensities, and the right graph shows the ratio of fluorescence intensities measured in the bleached and unbleached regions (bleached/unbleached). **(C)** Table showing the time required for the bleached/unbleached intensity ratio to reach 0.8 (80% equilibration) in 10 independent photobleaching experiments. Numerical data used to generate the graph are provided in Source Data 1. Scale bars are 10 µm.

Together, these results suggest that Sm contributes to the formation and maintenance of a distinct environment along the Y-loops, within which nascent transcripts are organized. Rather than directly binding to nascent transcripts, these proteins appear to act as structural or organizational components of the compartment, potentially creating a local environment that supports the spatial organization of nascent RNA for efficient elongation and maturation.

### *kl-3* processing occurs near the nucleolus-like body

Our findings suggest that nascent transcripts maintain a distinct spatial organization along the distal regions of the Y-loop, potentially undergoing the progression of elongation of the exceptionally large *kl-3* intron5. We next sought to determine how transcription initiation, elongation, and splicing-associated events are spatially organized for this same gene.

Previous work by Fingerhut et al. proposed that the nucleolus-like body (NLB) serves as a site of RNA polymerase II activation during Y-loop transcription^21^. To assess whether these events occur in distinct sites on the Y-loops, we examined the distribution of histone modifications associated with different events.

We first examined H3K4me3, a histone modification associated with active promoters and transcriptional initiation ^34^. H3K4me3 was enriched near the NLB (Figure 5A), consistent with the proposed role of this region in transcriptional activation and initiation ^21^. This localization suggests that the proximal region of the Y-loop adjacent to the NLB may represent a specialized transcriptional domain where promoter activation and the initiation of RNA polymerase II activity occur.

**Figure 5.**
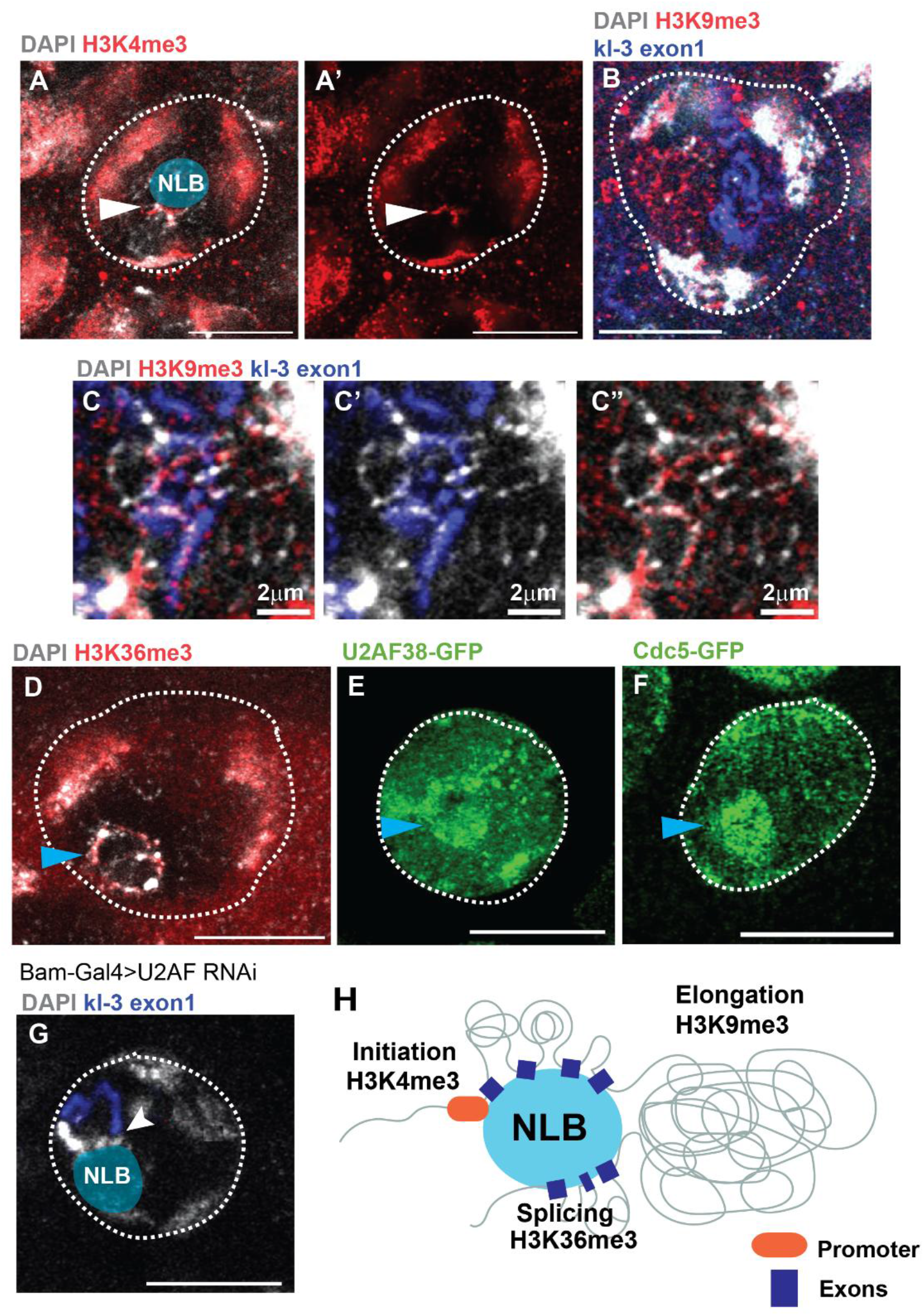
**(A)** Representative immunofluorescence image of a mid-stage SC nucleus from *w¹¹¹⁸* flies stained with an antibody against H3K4me3 (red). White dotted lines outline the SC nucleus. Arrowheads indicate H3K4me3-positive chromatin associated with the NLB. DAPI (gray). **(B, C)** Representative images of mid-stage SC nuclei from *w¹¹¹⁸* flies subjected to combined immunofluorescence staining and RNA FISH using an antibody against H3K9me3 and a probe against *kl-3* exon 1 (red and blue, respectively). White dotted lines outline SC nuclei. **(D)** Representative immunofluorescence image of a mid-stage SC nucleus from *w¹¹¹⁸* flies stained with an antibody against H3K36me3 (red). White dotted lines outline the SC nucleus. The blue arrowhead indicates the NLB. **(E, F)** Representative images of mid-stage SC nuclei from U2af38-GFP (E) or Cdc5-GFP (F) flies. Live samples were used. White dotted lines outline SC nuclei. Blue arrowheads indicate the NLB. **(G)** Representative RNA FISH image of a mid-stage SC nucleus from testes of *bamGal4*-driven *U2af38 RNAi* flies, hybridized with a probe against *kl-3* exon 1. The white arrowhead indicates the position where the *kl-3* signal is interrupted at the NLB. **(H)** Schematic interpretation of *kl-3* Y-loop organization. DAPI (gray). Blue circles indicate the approximate locations of the NLBs. Scale bars are 10 µm unless otherwise indicated in the images.

We next examined H3K9me3. While H3K9me3 is known to be a repressive heterochromatic mark, it is often enriched in the active genes located in constitutive heterochromatin including *kl-3* and other Y-chromosome genes ^35^, and is required for their expression ^36–39^. In contrast to the restricted distribution of H3K4me3, H3K9me3 was detected throughout the distal Y-loop regions, colocalizing with kl-3 exon1 RNA FISH (Figure 5B, C). This distribution is consistent with the presence of transcriptionally active polymerase along the extended Y-loop and suggests that the chromatin environment changes as transcription progresses from the proximal region toward distal loop regions.

Surprisingly, H3K36me3, a histone modification associated with splicing^40,41^, was enriched near the NLB (Figure 5D). This localization was unexpected given its established association with splicing regulation. Its enrichment near the NLB raises the possibility that RNA-processing events are spatially concentrated in this region.

To further investigate this possibility, we examined the localization of splicing-associated factors. We found that U2af38, a component involved in recognition of the intron 3’ splice site and early spliceosome assembly ^42^, localized near the NLB (Figure 5E). Similarly, Cdc5, another component of the spliceosome complex ^43^, was detected in the same region (Figure 5F). The localization of these splicing-associated factors near the NLB suggests that the splicing event is occurring in this region. Consistently, we often observed interruption of *kl-3* transcript at the NLB in *U2af38* RNAi (Figure 5G), indicating that the transcript is paused due to the splicing defect and following checkpoint activation at this region^44^.

Intriguingly, Fingerhut et al.^21^ reported the enrichment of intron-exon junction-containing transcripts near the NLB. Together with our observations of H3K36me3, U2AF, and CDC5 localization in this region, these findings support the possibility that pre-mRNA splicing occurs near the NLB. In particular, the convergence of splicing-associated factors and intron-exon junction-containing transcripts near the NLB suggests that this region may recruit exon-intron junction and serve as a site of co-transcriptional RNA processing (Figure 5H).

Based on these findings, we propose that Y-loop transcription and RNA processing are spatially organized into distinct functional domains. Transcriptional initiation may occur near the NLB, followed by elongation along distal Y-loop regions, while intronic and exon-containing nascent transcripts are organized in a manner that facilitates their subsequent recruitment to the NLB-associated RNA-processing machinery. This model provides a potential explanation for how an exceptionally long transcription unit can maintain efficient RNA processing despite its extended chromosomal organization.

## Discussion

Our findings reveal a spatially organized mechanism for the expression of the exceptionally long *kl-3* gene on the *Drosophila* Y chromosome. We propose a model in which nascent transcripts undergo distinct organizational and processing steps during transcription. In this model, exon-containing nascent RNA remains tightly associated with the DNA template strand, which has previously been suggested by structural studies ^3,5^, but has never been visualized in vivo. In addition, we show intronic RNA sequences form a compact outer layer surrounding the exon-associated RNA. Sm contributes to maintaining such nascent transcript organization, which is required for efficient elongation and maturation of *kl-3* transcripts.

Transcriptional activity is spatially distributed along the Y-loop, with transcriptional initiation and promoter-associated chromatin features enriched near the NLB, while active polymerase and intronic transcripts are detected throughout distal loop regions. The distinct distributions of H3K4me3, H3K9me3, and H3K36me3 further suggest that the chromatin environment changes along the transcription unit in association with different stages of transcription and RNA processing. We propose that the NLB serves as a specialized RNA-processing domain where nascent transcripts and splicing-associated factors, including U2af38 and Cdc5, are concentrated. The spatial organization of exon- and intron-containing sequences may facilitate the recruitment or presentation of nascent transcripts to this processing machinery.

Thus, our model suggests that the spatial organization of nascent RNA, together with the compartmentalization of transcriptional and RNA-processing activities, provides a mechanism for coordinating the expression of exceptionally long Y-linked genes. More broadly, this principle may represent a general strategy for organizing gene expression, in which the spatial arrangement of nascent transcripts and the establishment of specialized nuclear environments facilitate the coordination of transcription, RNA processing, and transcript maturation. Such spatial organization may be particularly relevant for genes with complex intron structures, extended transcription units, or high demands for RNA processing, suggesting that nuclear compartmentalization contributes to the regulation of gene expression across diverse genomic contexts.

## Methods

### Fly husbandry and strains

Flies were raised on standard Bloomington medium (Lab Express) at 25°C and young flies (0- to 7-day-old adults) were used for all experiments. Control flies were either the parental Bam-GAL4 stock or a sibling from the same genetic cross. The following fly stocks were obtained from the Bloomington *Drosophila* Stock Center (BDSC); *w1118* (BDSC3605), *UAS-U2af38RNAi^TRiP.HMS04505^* (BDSC57585), *BamGal4/X* (BDSC80579)*, Df(3L)BSC371* (Blanks deficiency, BDSC24395), *blanks^KG00084^* (BDSC13914), *UAS-smRNAi*^TRiP.HMC05543^ (BDSC64524), *sm^[MI04033-GFSTF.1]^* (BDSC59815), *Cdc5-GFP.FPTB* (BDSC66585), *UAS-Prp5RNAi^TRiP.HMC03786^* (BDSC55638), *UAS-Prp5RNAi^TRiP.HMJ24050^* (BDSC62507). The following fly stocks were obtained from the Vienna *Drosophila* Resource Center: *U2af38-GFP* (v318694). *GFP-blanks* transgenic line (GFP-tagged Blanks expressed from endogenous promoter) was a gift from Yukiko Yamashita^18^.

### Stellaris RNA FISH

Stellaris RNA FISH was performed as described previously^45^ with slight modifications. Testes were dissected in 1X phosphate buffered saline (PBS) and then fixed in 1 ml of 4% formaldehyde/PBS for 45 minutes. Fixed testes were rinsed 2 times with 1 ml of 1X PBS, then resuspended in 1 ml of chilled 70% ethanol, and incubated for 1 hour – overnight at 4 °C. Testes were rinsed briefly in 1 ml of wash buffer (2X SSC (ThermoFisher) and 10% deionized formamide), then incubated overnight at 37 °C in the dark with 50 nM of probe in 100 µl of Hybridization Buffer containing 2X SSC, 10% dextran sulfate (MilliporeSigma), 1 μg/μl of yeast tRNA (MilliporeSigma), 2 mM vanadyl ribonucleoside complex (NEB), 0.02% BSA (ThermoFisher), and 10% deionized formamide. Then, testes were washed 2 times for 30 minutes each at 37 °C in the dark in 1 ml of prewarmed wash buffer (2X SSC, 10% formamide) and resuspended in a drop of VECTASHIELD with 4’,6-diamidino-2-phenylindole (DAPI) (Vector Lab).

Quasar 570 labeled Stellaris probe against *Juan* and Quasar 670 labeled Stellaris probes against *HMS Beagle* were designed with LGC Biosearch Technologies (www.biosearchtech.com/stellarisdesigner). Quasar 670 labeled Stellaris probe against *kl-3* exon 1 and Quasar 570 labeled Stellaris probe against *kl-5* exons 1-6 was obtained from LGC Biosearch Technologies using probe sequences described previously^18^. The AATAC satellite probe was obtained from Integrated DNA Technologies. Probe and target sequences are provided in Spreadsheet1.

### Hybridization chain reaction (HCR) HiFi RNA FISH

HCR RNA FISH was performed as described previously^46^ with slight modifications. Testes were dissected in 1X PBS and then fixed in 1 ml of 4% formaldehyde/PBS for 30 minutes. Fixed testes were washed for five minutes each in 1 ml 1X PBS + 0.2% Triton X-100 (Thermo Fisher) (PBST), 1 ml 50/50 PBST/5X SSC + 0.1% Tween-20 (5XSSCT), 1ml 5XSSCT. Testes were prehybridized in 200 μl Hybridization Buffer (Molecular Instruments) at 37°C for 30 minutes. Testes were then incubated overnight at 37 °C in the dark with 20 nM of probe in 200 µl of Hybridization Buffer. Testes were washed 15 minutes in 1 ml 100% Probe Wash Buffer (Molecular Instruments), then 5 minutes each in 1ml 75% Probe Wash Buffer/25% 5XSSCT, 1ml 50% Probe Wash Buffer/50% 5XSSCT, and 25% Probe Wash Buffer/75% 5XSSCT, all at 37°C. Testes were prehybridized 5 minutes with 200 μl HCR Gold Amplification Buffer (Molecular Instruments) at room temperature. During the wash steps, hairpins were prepared by heating in a PCR machine at 95°C for 90 seconds, then cooling at room temperature in the dark for 30 minutes. Testes were then incubated at room temperature 6 hours – overnight in 200 μl Amplification Buffer with 6 pmol hairpins. After incubation, testes were washed four times for 15 minutes each with HCR Gold Amplifier Wash Buffer (Molecular Instruments) before resuspension in a drop of VECTASHIELD with DAPI.

HCR Gold probes against *kl-3* exon 5-exon 6 and *kl-3* exon5-intron5 were designed by Molecular Instruments. Probe sequences can be found in Spreadsheet1.

To combine HCR FISH with smFISH, the HCR FISH protocol was used. smFISH probes were added during the probe hybridization step at the same concentration used for smFISH. Images were taken using a Zeiss LSM800 confocal microscope with airyscan module with 63X oil immersion objective (NA = 1.4) using Zen software. Images were processed by image J/FIJI.

### Larval brain squash DNA fluorescence in situ hybridization

We adapted the DNA FISH protocol for mitotic neuroblast from Larracuente & Ferree, 2015^47^ with slight modifications. Brains from male third instar larvae were dissected in 1X PBS and moved to a Superfrost Plus slide. Excess PBS was removed and replaced with 25µl of 0.5% sodium citrate. The sodium citrate was removed and replaced with a fresh 25µl of 0.5% sodium citrate. The sample was then incubated for 10 minutes at room temperature. The sodium citrate was removed and replaced with 25µl fixative solution (4% formaldehyde in PBS: glacial acetic acid [55%: 45%]). The fixative solution was removed and replaced with a fresh 25µl of fixative solution. The sample was then incubated for 4 minutes at room temperature. A coverslip was placed on the slide, and samples were squashed by thumb for 20 seconds. The slide was then submerged in liquid nitrogen until boiling stopped. The coverslip was quickly removed using a razor blade, and the slide was washed in 100% ethanol for 5 minutes at room temperature. The slide was removed from ethanol and dried before adding enough 50% Formamide in 2X SSCT (2X SSC + 0.1% Tween-20) to cover the sample. A coverslip was placed on the slide before heating the sample at 92°C for 30 minutes. Slides were cooled on ice for at least 5 minutes before removing the coverslip and adding hybridization mix. The hybridization mix contained 50% formamide and 10% dextran sulfate in 2X SSCT with 25pmol of probe and 1.25µl of RNase Cocktail Enzyme Mix (Thermo-Fisher). Water was added up to a total volume of 100µl. The hybridization mix (100 µl/sample) was denatured at 65 °C for 5 min and kept on ice before adding to the samples. A fresh coverslip was placed on the slide before incubating at 37°C overnight in a dark, humid chamber. The coverslip was then removed, and the slide was washed three times: (1) 30 minutes in 50% Formamide in 2X SSCT at 37°C, (2) 10 minutes in 20% Formamide in 2X SSCT at room temperature, (3) 10 minutes in 2X SSCT at room temperature. A drop of VECTASHIELD with DAPI was added to the sample and a fresh coverslip was applied. Probe sequences are listed in Spreadsheet1. Images were taken using a Zeiss LSM800 confocal microscope with airyscan module with 63X oil immersion objective (NA = 1.4) using Zen software. Images were processed by image J/FIJI.

### Immunofluorescence staining

Testes were dissected in PBS and fixed in 4% paraformaldehyde/PBS for 30 min. Samples were then washed in PBST (PBS + 0.2% TritonX-100) for 60 minutes (20 minutes x3), followed by overnight incubation with primary antibody in PBST containing 3% bovine serum albumin (BSA) at 4°C. Samples were then washed for 60 minutes in PBST, incubated with secondary antibody in 3% BSA in PBST at room temperature for 2 hours and then washed for 60 minutes (three times for 20 minutes each) in PBST. Samples were then mounted using VECTASHIELD with DAPI. The primary antibodies used were as follows: Rabbit anti H3K4me3 (1:100, Catalog#: 39916); Rabbit anti H3K9me3 (1:100, Catalog#: 39065); Rabbit anti Pol2 pSer5 (1:100, Catalog#:39234); Mouse anti H3K36me3 (1:100, Catalog#: 91266) were obtained from Active Motif. AlexaFluor-conjugated secondary antibodies (Abcam) were used at a 1:200 dilution. Images were acquired on a Zeiss LSM800 airyscan with a 63X/1.4 NA oil objective. Images were processed by image J/FIJI.

### Immunofluorescence staining combined with Stellaris RNA FISH

After Stellaris RNA FISH procedure, samples were fixed in 4% paraformaldehyde/PBS for 10 min, then samples washed in PBST (PBS + 0.2% TritonX-100) for 60 minutes (20 minutes x3), then followed by primary antibody incubation and later regular procedures of immunofluorescence staining (see above).

### Line scan of RNA FISH images

Image-J/Fiji was used for image quantification. Z-stack series were taken using a 3X/1.4 NA oil objective on a Zeiss LSM800 confocal microscope with airyscan module. Images were Airyscan processed with standard strength. RNA FISH intensity was measured in SC nuclei using a single slice from a z-stack. The line tool was used to draw a line perpendicular across a loop signal and intensity was measured in individual channels across the line using the Plot Profile tools. Intensity was then compared between channels to determine signal localization.

### Short-term live imaging and fluorescence recovery after photobleaching (FRAP)

Testes from newly eclosed flies were dissected into Schneider’s *Drosophila* medium containing 10% fetal bovine serum. These testes were placed onto Gold Seal Rite-On Micro Slides’ 2 etched rings with media, then covered with coverslips. Images were immediately taken using a Zeiss LSM800 confocal microscope with a 63× oil immersion objective (NA = 1.4).

FRAP imaging and analysis were performed as described previously^48^. Signal recovery was undertaken using a Zeiss LSM800 with airyscan module by using 1AU-pinhole with 63X oil immersion objective (NA = 1.4). Zen software was used for programming each experiment. Encircled areas of interest (randomly chosen 6µm-diameter circles covering the Blanks-GFP-enriched region of a spermatocyte nucleus) were photobleached using the 488 nm laser (laser power; 100%, iterations; 5). Recovery images were taken every 2 seconds over 60 seconds for single z-plane. Images were processed by image J/FIJI.

### Statistical analysis and graphing

No statistical methods were used to predetermine sample size. The experiments were not randomized. The investigators were not blinded to allocation during experiments and outcome assessment. All experiments were independently repeated at least three times to confirm the results. Statistical analysis and graphing were performed using GraphPad Prism 10 software. Individual numerical values displayed in a graph are provided in Source data1.

## Acknowledgements

We thank Yukiko Yamashita, the Vienna *Drosophila* Resource Center, and the Bloomington *Drosophila* Stock Center for reagents. This research is supported by R35GM128678 from the National Institute for General Medical Sciences and a start-up fund from UConn Health (to M.I.).

---

**Figure S1.**
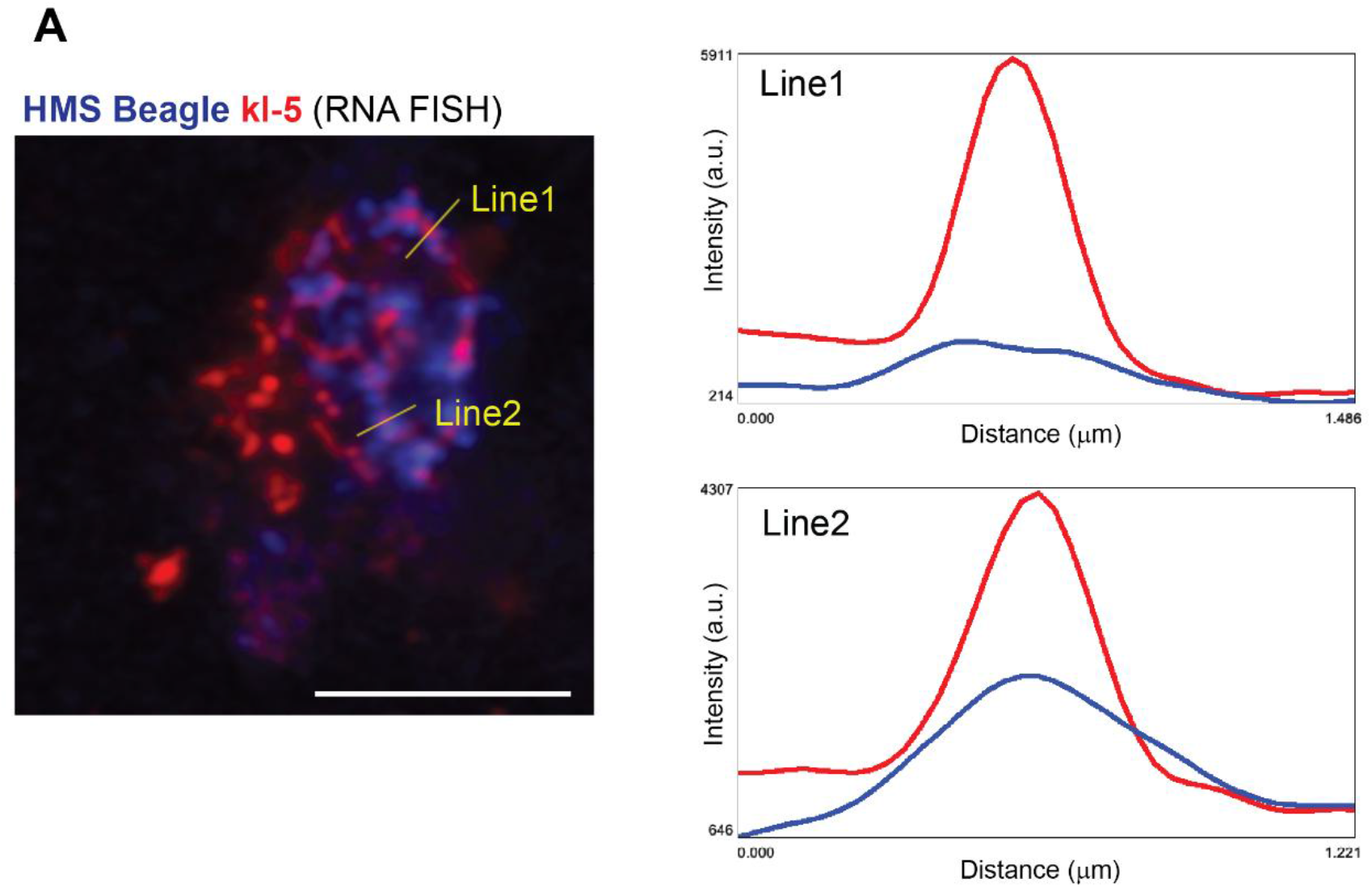
**(A)** Representative RNA FISH image of an SC nucleus from *w¹¹¹⁸* flies. The probes used are shown in the corresponding colors. The yellow lines indicate the regions used for the line scans shown on the right. Scale bar: 10 µm.

## Movie S1

Representative time-lapse movie corresponding to Figure 4A and B, showing Sm-GFP fluorescence before photobleaching, immediately after photobleaching, and during fluorescence recovery. Scale bar: 10 µm.

## Spreadsheet 1

Sequence information for the FISH probes used in this study.

## Source Data 1

Numerical values underlying all graphs and plots presented in the figures.

